# Circadian redox rhythm gates immune-induced cell death distinctly from the genetic clock

**DOI:** 10.1101/2023.04.21.535069

**Authors:** Sargis Karapetyan, Musoki Mwimba, Xinnian Dong

## Abstract

Organisms use circadian clocks to synchronize physiological processes to anticipate the Earth’s day-night cycles and regulate responses to environmental stresses to gain competitive advantage^1^. While divergent genetic clocks have been studied extensively in bacteria, fungi, plants, and animals, a conserved circadian redox rhythm has only recently been reported and hypothesized to be a more ancient clock^2, 3^. However, it is controversial whether the redox rhythm serves as an independent clock and controls specific biological processes^4^. Here, we uncovered the coexistence of redox and genetic rhythms with distinct period lengths and transcriptional targets through concurrent metabolic and transcriptional time-course measurements in an *Arabidopsis* long-period clock mutant^5^. Analysis of the target genes indicated regulation of the immune-induced programmed cell death (PCD) by the redox rhythm. Moreover, this time-of-day-sensitive PCD was eliminated by redox perturbation and by blocking the signalling pathway of the plant defence hormones jasmonic acid/ethylene, while remaining intact in a genetic-clock-impaired line. We demonstrate that compared to robust genetic clocks, the more sensitive circadian redox rhythm serves as a signalling hub in regulating incidental energy-intensive processes, such as immune-induced PCD^6^, to provide organisms a flexible strategy to prevent metabolic overload caused by stress, a unique role for the redox oscillator.

---

Rhythmic daily behaviours in plants and animals have been observed since antiquity^7^. However, the mechanistic basis of circadian clocks was not revealed until the 20^th^ century, with the discoveries of the first clock genes^8–10^. These genes encode an internal genetic oscillator consisting of transcription-translation feedback loops (TTFL), which synchronizes the physiological processes/behaviours with diurnal environmental changes and “gates” (i.e., modulates the strength of) responses to biotic and abiotic changes to improve fitness of the organism^1, 11^. In addition to the genetic clock, recent discoveries of circadian oscillations of the peroxiredoxin oxidation state in anucleate red blood cells and in mutants or lines that lack oscillations of the canonical TTFL circuitry in the absence of known entrainment signals (called in this manuscript “clock-dead” for simplicity)^2, 3^, together with the detection of a significant number of oscillating transcripts in these mutants^12, 13^, suggest the existence of a parallel metabolic clock. Furthermore, the conservation of this redox rhythm across all lineages of life indicates its possible importance in evolution since the great oxidation event^3^. However, after the initial excitement of these ground-breaking discoveries, concerns over the observed oscillations in clock-dead mutants being driven by unknown external factors^4^, along with the lack of a clear physiological output regulated specifically by the circadian redox rhythm, have cast doubt on the autonomy and the significance of the redox oscillator. Therefore, while an interplay between the genetic clock and the redox rhythm has been well established^14–16^, the biological significance of the redox oscillation itself remains largely unknown.

## The genetic clock and redox rhythm are disentangled in a long-period clock mutant

The difficulty in studying the distinct circadian function of the redox rhythm stems from the fact that it is intricately intertwined with the genetic clock. Disruption of the redox rhythm can affect the period, phase or the amplitude of the genetic clocks depending on their molecular architectures and the nature of the disruption^14, 15^ and vice versa^3^. Thus, genetic-clock-dead mutants have been used in attempt to identify the specific transcriptional targets of the redox rhythm. However, the scant overlap between the oscillating transcripts in the clock-dead mutants and the wild type (WT)^12, 13^ raises the concern that the targets identified in the mutants might not be functionally relevant for the WT. To disentangle the genetic and redox rhythms, we decided to use a period-length mutant to simultaneously identify distinct output genes of both based on their oscillation periods. We used the *Arabidopsis* mutant of the clock genes *PSEUDO-RESPONSE REGULATORs (PRRs) 7* and *9* (*prr7 prr9*) which has the intrinsic genetic clock periods of approximately 24 h at 12 °C, 32 h at 22 °C (typical temperature for *Arabidopsis* experiments) and 36 h at 30 °C^5^. Therefore, in contrast to the WT, which has the ability to maintain the ∼24 h period across a physiological range of temperatures, the *prr7 prr9* mutant is defective in temperature compensation. This temperature-dependent period variation in the *prr7 prr9* mutant makes it a tractable system to separate the redox and genetic rhythms based on oscillation periods.

We grew WT and *prr7 prr9* plants in temperature- and humidity-controlled (22 °C, 65% RH) Percival chambers^17^, under 12h light/12 h dark (LD) conditions to first entrain both the genetic clock and the redox rhythm by the light/dark cycles. To examine the intrinsic behaviours of the two clocks, we released the plants to constant light (LL) conditions at either 22 °C or 30 °C and collected plant tissue for concurrent metabolic and transcriptomic analyses from 24 h to 92 h in LL (Fig. 1a). Since glutathione plays a central role in maintaining cellular redox homeostasis in plants^18^, we used the reduced (GSH) and oxidized (GSSG) glutathione pair to represent the redox rhythm. For transcriptomic measurements, we used the multiplex RNA annealing selection ligation-sequencing (RASL-seq) approach^19^ to analyse the expression of a selected pool of ∼700 genes as an alternative to RNA-seq to accommodate the large sample number for the time-course experiments (two genotypes, two temperatures and 18 time points with 6 replicates for 22°C and 3 replicates for 30 °C).

**Fig. 1.**
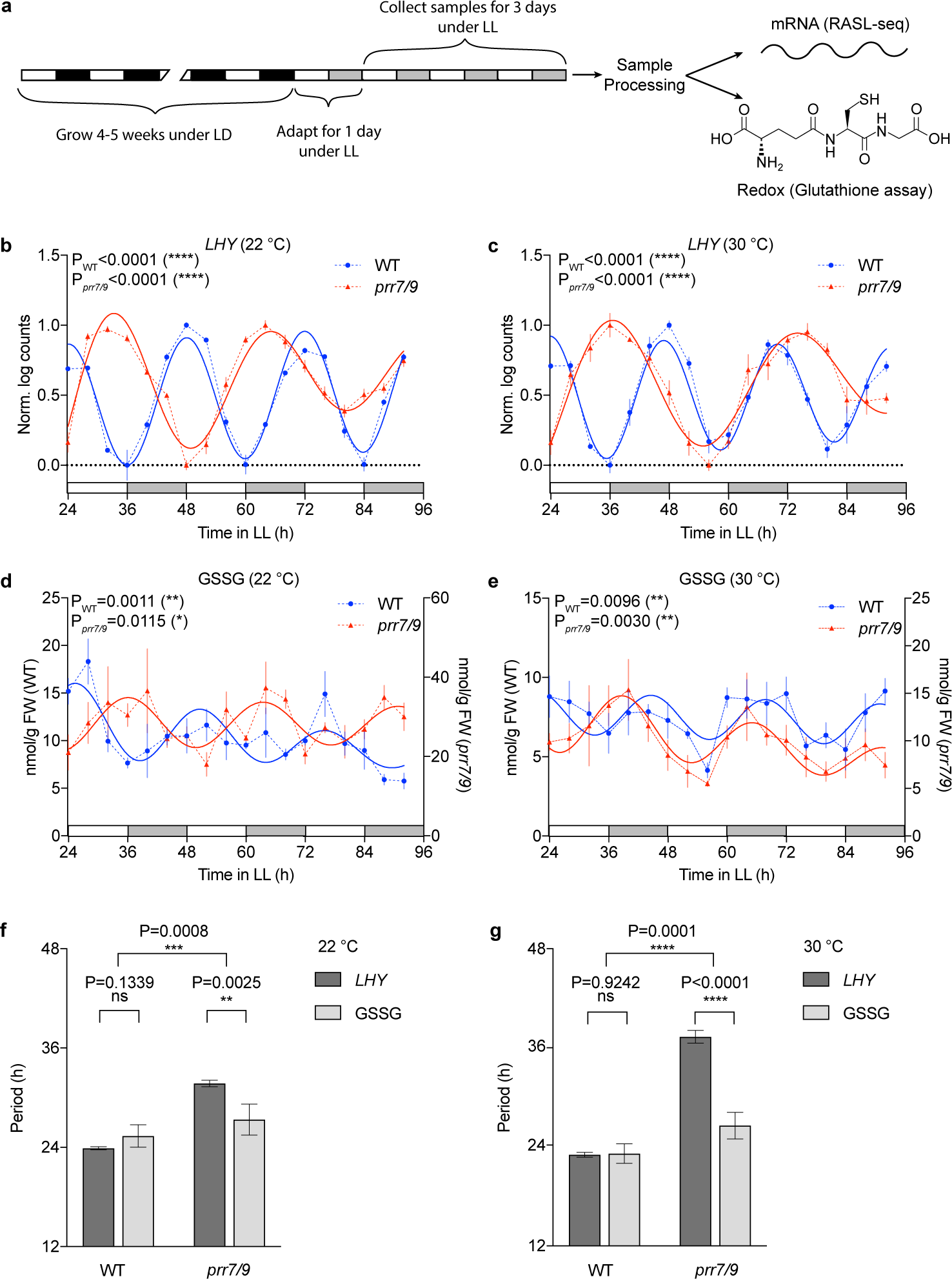
The redox rhythm and genetic clock are separated in the *prr7 prr9* double mutant. **a**, Workflow of the sample collection and processing for transcriptional and metabolic measurements. WT and *prr7 prr9* plants were first grown under light/dark (LD)-entraining conditions for 4-5 weeks and then released to constant light (LL) conditions. After 24 h of adaptation, samples were collected concurrently for mRNA (RASL-seq) and glutathione (GSH/GSSG) measurements. **b**, **c**, Normalized *LHY* transcript levels in WT and *prr7 prr9* at 22 °C (**b**) and 30 °C (**c**) starting from 24 h in LL. Solid lines represent the harmonic regression, indicating statistically significant oscillations. n ≥ 5 (**b**) and n = 3 (**c**). **d**, **e**, Oxidized glutathione (GSSG) levels in WT and *prr7 prr9* at 22 °C (**d**) and 30 °C (**e**) measured starting from 24 h after being moved to LL. Solid lines represent the harmonic regression, indicating statistically significant oscillations. n = 3 (**d**) and n ≥ 4 (**e**). **f**, **g**, Estimated periods from the harmonic regression for **b**-**e**. Individual values were compared using Student’s t-test, while the interaction was tested using 2-way ANOVA; ns, not significant. All values are means ± SEM.

Harmonic regression analysis of the time-course data^14^, revealed that in contrast to the core genetic clock gene *LATE ELONGATED HYPOCOTYL (LHY)* which showed longer periods in the *prr7 prr9* mutant than in the WT (Fig. 1b, c), GSSG exhibited close-to-WT periods in the *prr7 prr9* mutant, albeit with different phases, at both 22 °C and 30 °C (Fig. 1d, e). Thus, the redox rhythm, represented by GSSG, is separated from the genetic clock based on the period lengths in the *prr7 prr9* mutant (Fig. 1f, g). Furthermore, while the elevated temperature increased the period of *LHY* from 31.71 h to 37.34 h in the *prr7 prr9* mutant, recapitulating the previously observed temperature overcompensation^5^, period of GSSG changed only slightly from 27.34 h to 26.57 h. Therefore, GSSG oscillation also appears to exhibit temperature compensation, a known hallmark of circadian clocks^20^, separate from the genetic clock.

Surprisingly, in contrast to GSSG, the total glutathione (GSSG + GSH) oscillation was tied to the genetic clock, likely through the regulation of the *GLUTAMATE-CYSTEINE LIGASE* (*GSH1*), which encodes the enzyme for catalysing the rate-limiting step of glutathione biosynthesis in *Arabidopsis*^21^ (Extended Data Fig. 1). Since the total glutathione and GSSG levels were measured in the same samples, the separation of GSH+GSSG and GSSG oscillations further supports the independence of the redox rhythm from the genetic clock. Thus, GSSG, composing a small fraction of the total glutathione pool, acts as a representation or a “sensor” for the GSH oxidation reactions constituting a branch of the redox rhythm. In contrast, the overall glutathione production is regulated by the genetic clock, providing a possible mechanism for the genetic clock to entrain the redox rhythm.

## Transcriptional targets of the genetic clock and the redox rhythm are distinguished based on periods

The different oscillation periods for the redox rhythm and the genetic clock in the *prr7 prr9* mutant provided the opportunity to identify their specific transcriptional targets. Due to significant differences in various period detection/estimation algorithms^22^, we decided to cluster the transcripts according to their normalized temporal signature instead of period. Therefore, we performed k-means clustering of the circadian time course RASL-seq data from 678 probes in the *prr7 prr9* mutant at 22 °C. After testing different k values, we chose k =16 to accommodate all possible clusters for the genes oscillating with the genetic clock period of approximately 32 h (8 points for each of the possible phases given by the time resolution of 4 h, multiplied by 2 to account for positive and negative skew). For genes in each cluster defined using the *prr7 prr9* at 22 °C data, we then identified those that are oscillatory in WT and the *prr7 prr9* mutant at 22 °C and 30 °C and calculated their periods using Biodare’s FFT-NLLS algorithm^22^ in these genetic backgrounds and temperatures. While a close-to-24 h period was observed across all clusters in the WT at 22 °C, larger period length variations were found in the *prr7 prr9* mutant with majority of the gene clusters oscillating with the expected genetic clock period of ∼32 h (Fig. 2a, b). However, clusters 2 and 6 (C2 and C6) contained multiple oscillatory genes (shown as dots) with periods in *prr7 prr9* close to that in WT. Since the output of k-means clustering can vary depending on the initial seed and the number of clusters, we also used a deterministic hierarchical algorithm to analyse the RASL-seq data, which largely recapitulated the results of the k-means clustering (Extended Data Fig. 2). Remarkably, oscillations for genes in both C2 and C6 also showed altered phase in *prr7 prr9* compared to WT, resembling that of GSSG (Figs. 1d, 2c, d), consistent with these genes being regulated by the redox rhythm. Surprisingly, at 30 °C, virtually none of the genes in C2 and C6 kept the shorter period (Extended Data Fig. 3). At this elevated temperature, these genes appeared to be driven by the genetic clock. Therefore, the C2 and C6 genes can be regulated by both the redox rhythm and the genetic clock, with the redox rhythm clock playing the primary role under the normal temperature and the genetic clock dominating at elevated temperatures. Among the genes studied in our RASL-seq, a single gene, *FLOWERING LOCUS T* (*FT*), reliably oscillated with the shorter periods under both temperatures (Extended Data Fig. 4a, b), in agreement with the near WT redox rhythm period observed in the *prr7 prr9* mutant (Fig. 1d, e). These results suggest that at 30 °C, while the C2 and C6 genes are delinked from the redox rhythm, *FT,* and perhaps other flowering time genes not present in our RASL-seq gene pool, is still under the circadian redox rhythm regulation.

**Fig. 2.**
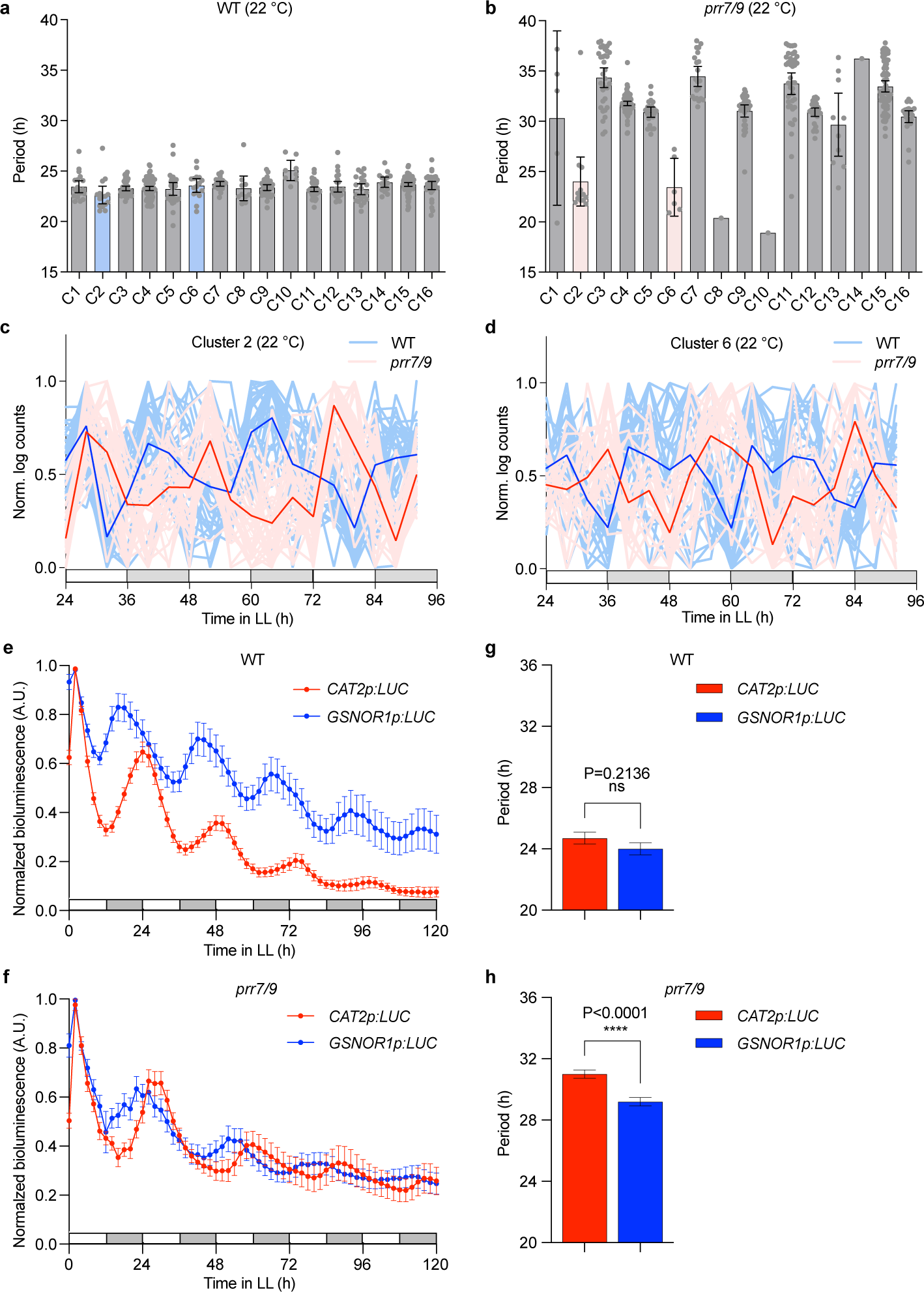
In *prr7 prr9*, two gene clusters oscillate with periods distinct from those of the genetic clock. **a**, **b**, Periods of oscillatory transcripts in WT (**a**) and *prr7 prr9* (**b**) at 22 °C. Clusters were generated based on k-means analysis of expression signatures of all transcripts in the RASL-seq dataset from the *prr7 prr9* at 22 °C samples. For each cluster, periods in WT and *prr7 prr9* were estimated using FFT-NLLS, with each dot representing the period of a single transcript. Clusters of interest (C2 and C6) are coloured light blue (**a**) and light red (**b**). **c**, **d**, Normalized expression levels of all genes in C2 (**c**) and C6 (**d**) in WT and *prr7 prr9* at 22 °C. Dark lines represent the means of each cluster, while light lines represent the expression pattern of the means of individual genes in the clusters. **e, f,** Normalized bioluminescence traces for independent T1 transformants of *CAT2p:LUC* and *GSNOR1p:LUC* in WT with n = 16 and n = 8, respectively (**e**) or in *prr7 prr9* with n = 13 and n = 10, respectively (**f**). **g, h,** Period estimation of independent T1 transformants from three separate experiments, combined using a linear mixed-effect model^59^, using FFT-NLLS for *CAT2p:LUC* and *GSNOR1p:LUC* in WT with n = 18 and n = 18, respectively (**g**) and in *prr7 prr9* with n = 24 and n =28, respectively (**h**). Only oscillations with RAE ≤ 0.7 were considered for the analysis. Periods were compared using Student’s t-test; ns, not significant. All values are means ± SEM.

Since time-course experiments may lead to inaccurate period estimation due to the low time resolution and plant sample variations, we next constructed bioluminescence reporter lines to verify our findings. Based on previous publications^14, 17^, we chose the promoter of *CATALASE 2* (*CAT2*) to drive the luciferase reporter as a known morning-phased output of the genetic clock, which, as expected, oscillated with a long period in *prr7 prr9* in our RASL-seq experiment (Extended Data Fig. 4c, d). To make a reporter for the redox rhythm output, we chose, from C6, the promoter of *S-NITROSOGLUTATHIONE REDUCTASE 1* (*GSNOR1*), which is the key regulator of S-nitrosothiol (SNO) levels in plants^23^. Besides its robust oscillation detected in our RASL-seq data (Extended Data Fig. 4e, f), the dependence of GSNOR1 activity on NADH and GSH^24^ further connects the gene to the redox rhythm. Both *CAT2p:LUC* and *GSNOR1p:LUC* constructs were transformed into WT and the *prr7 prr9* backgrounds and multiple independent T1 lines were then assessed for period length to avoid insertion-position effects on the reporter expression. In agreement with the RASL-seq data, significant differences in period lengths of *CAT2p:LUC* and *GSNOR1p:LUC* oscillations were detected in *prr7 prr9,* but not in WT (Fig. 2e-h), after both were transferred from LD to LL, across three independent experiments, confirming that these genes are driven by distinct oscillators. Since the age of the leaf may affect the period of the circadian clock^25^ within the same plant, we measured the luminescence of the shoot apex meristem (SAM), whose circadian rhythms are thought to exert dominance over those of other tissues^26^. We observed, for the first time, the coexistence of two distinct circadian rhythms within the same tissue. To ascertain that the period separation is not limited this pair of genes, we tested another pair of reporters. *CATALASE 3* (*CAT3*) was chosen as an evening-phased genetic clock output^14^ to ensure that both morning- and evening-phased genetic clock outputs were separated from the redox rhythm. Simultaneously, the *GLUCOSE-6-PHOSPHATE DEHYDROGENASE 6* (*G6PD6*) gene encoding an enzyme from the oxidative pentose phosphate pathway (PPP)^27^ was chosen because PPP is a hub for the interplay between the redox rhythm and the genetic clock in mammals^15, 16^ and the *G6PD6* gene represents a different redox-regulated gene cluster (C2 in Fig. 2b). We found that *G6PD6p:LUC* oscillated with a significantly shorter period than *CAT3p:LUC* (Extended Data Fig. 5), further confirming the separation of the redox rhythm and the genetic clock in the *prr7 prr9* mutant.

## Redox rhythm regulates immune-induced programmed cell death

To reveal the biological functions of those redox-rhythm-regulated targets (i.e., clusters 2 and 6), gene ontology (GO) enrichment analysis was performed, with the entire RASL-seq probe collection used for clustering as the background. We found that, even in this probe set that contains mostly genes involved in different immune responses, the clusters were significantly enriched with those involved in the regulation of cell death (including immune-induced PCD, known as hypersensitive response in plants) and defence response (Fig. 3a). Further STRING (Search Tool for the Retrieval of Interacting Genes/Proteins) analysis uncovered a highly interconnected network, with known regulators of immune-induced cell death genes such as *SUPPRESSOR OF BIR1-1* (*SOBIR1*)^28^, *PHYTOALEXIN DEFICIENT 4* (*PAD4*) and *ENHANCED DISEASE SUSCEPTIBILITY 1* (*EDS1*)^29^ in hub positions (Fig. 3b). Moreover, consistent with our clustering analysis based on temporal expression signatures, STRING shows high degree of co-expression of genes in clusters 2 and 6 based on their publicly available data (black lines in Fig. 3b). Based on identities of these genes, we hypothesized that circadian redox rhythm regulates PCD during the effector-triggered immunity (ETI), which occurs when the presence of a pathogen effector is recognized by its cognate nucleotide-binding site, leucine-rich repeat (NB-LRR) receptor in the plant host and is associated with reactive oxygen species burst^30^ and transcriptional and translational programming fuelled by a significant increase in ATP levels^6, 31^. However, while the time-of-day sensitivity to ETI-triggered PCD has been reported previously^32^, no mechanism has been proposed for this circadian regulation. Remarkably, in agreement with the delinking of C2 and C6 genes from the redox rhythm at 30 °C (Extended Data Fig. 3), ETI-mediated PCD is well known to be inhibited at elevated temperatures^33^.

**Fig. 3.**
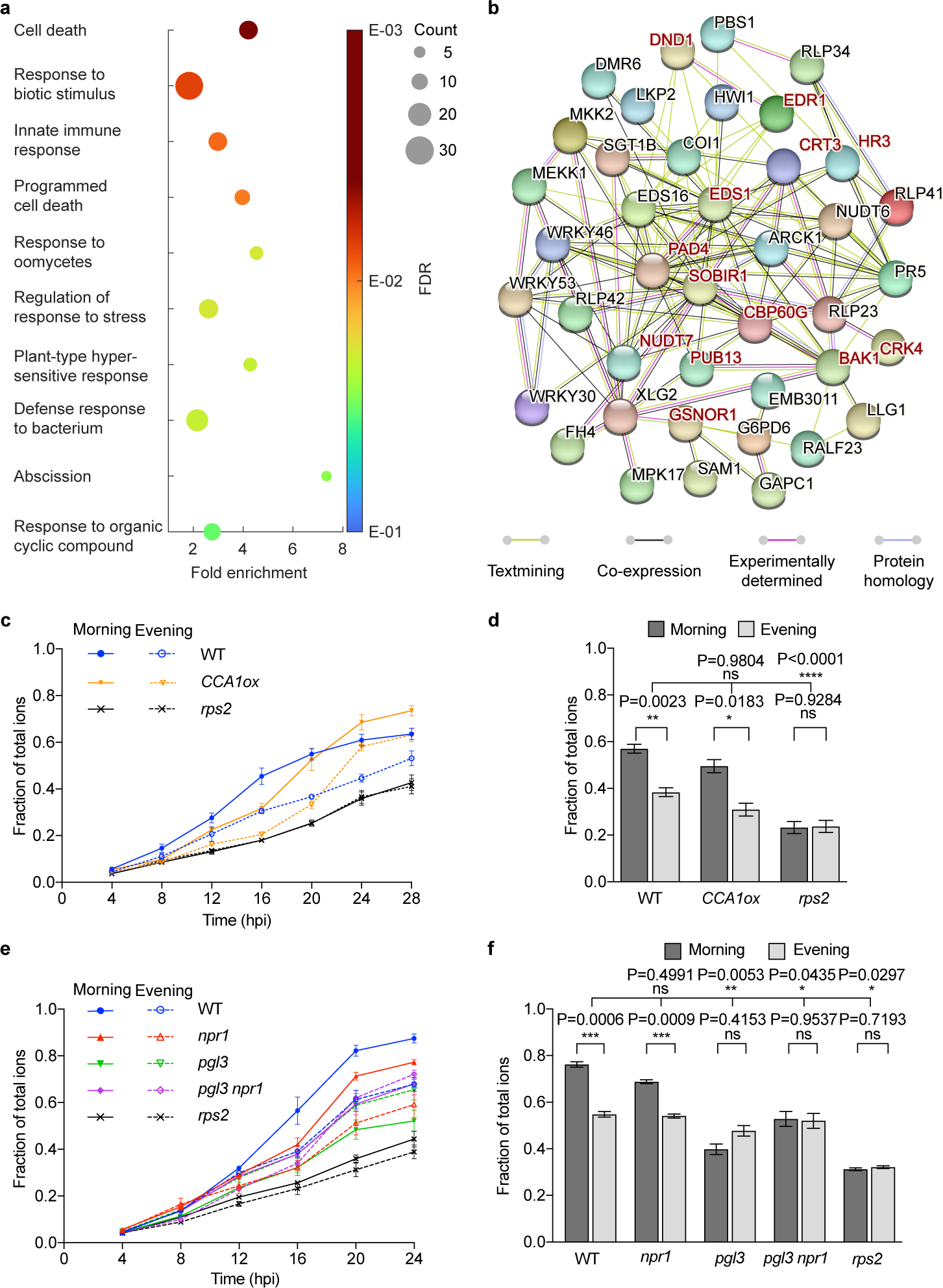
Redox rhythm regulates immune-induced PCD independent of the genetic clock. **a**, GO terms for combined C2 and C6 genes, sorted by the false discovery rate (FDR). The total RASL-seq gene pool was used as the background. **b**, StringDB analysis for C2 and C6 cluster genes using textmining, co-expression, experiments, and protein homology. Interaction score was set to 0.400. All genes belonging to the top GO term, cell death, are present and labelled in dark red letters. **c**, Time-course of ion leakage (a measure of cell death). After infiltration with *Psm* ES4326/*AvrRpt2* in WT, *CCA1ox*, and *rps2*, ion leakage was estimated by measuring conductivity of each timepoint normalized to conductivity of total ions for each sample. n = 3. **d**, Analysis of normalized conductivity at 20 hours post infiltration (hpi) from three separate experiments combined using a linear mixed-effect model. n = 9. Individual values were compared using Student’s t-test, while the interaction with the WT was tested using 2-way ANOVA; ns, not significant. **e**, After infiltration with *Psm* ES4326/*AvrRpt2* in WT, *npr1*, *pgl3*, *pgl3 npr1*, and *rps2,* ion leakage was estimated as in (**c**). n = 3. **f**, Normalized conductivity at 20 hpi from three separate experiments was analysed as in (**d**). n = 9. All values are means ± SEM.

To test the circadian control of PCD by the redox rhythm, we used bacterial pathogen *Pseudomonas syringae* pv. *maculicola* ES4326 carrying the effector gene *avrRpt2* (*Psm* ES4326/*avrRpt2*) to infect the WT, *CIRCADIAN CLOCK ASSOCIATED 1* overexpressing (*CCA1ox*) clock-dead line^34^ and the mutant lacking the NB-LRR receptor for avrRpt2, *resistant to p. syringae 2* (*rps2*). The infection was performed at subjective morning and evening, after 2 days of growth under constant light (LL) to eliminate any transient oscillations in *CCA1ox*. In agreement with our hypothesis, both WT and *CCA1ox* showed stronger ETI-associated PCD in the subjective morning than in the subjective evening (Fig. 3c, d), even though *CCA1ox* has been reported previously to lack the time-of-day-sensitive basal resistance to the bacterial pathogen without the avrRpt2 effector^35^. In contrast to *CCA1ox*, the mutant of *6-PHOSPHOGLUCONOLACTONASE 3* (*PGL3*), encoding a key enzyme in the oxidative section of the plastidic PPP, which has markedly altered glutathione levels and constitutively enhanced resistance^36^, had decreased PCD and lost the time-of-day sensitivity (Fig. 3e, f). This result is consistent with the finding that a functional PPP is required in the human red blood cells for maintaining the redox rhythm^37^. Since the basal resistance phenotype of *pgl3* is dependent on *NONEXPRESSOR OF PATHOGENESIS-RELATED GENES 1* (NPR1)^36^, shown to be the redox-rhythm-sensitive^14^, we also tested the *npr1* single and the *pgl3 npr1* double mutants for ETI-induced PCD. We found that the *npr1* single mutant still had time-of-day sensitivity to immune-triggered PCD, but when combined with *pgl3,* it could not rescue the PCD phenotype of *pgl3* (Fig. 3e, f), indicating that the lack of circadian PCD in *pgl3* is not due to constitutive activation of NPR1-mediated resistance.

## Redox rhythm regulates PCD through the jasmonic acid/ethylene defence pathway

The independence of the circadian PCD from NPR1, a key immune regulator required for salicylic acid (SA)-mediated promotion of cell survival against biotic and abiotic stresses^38^, suggests that the redox rhythm regulates ETI-associated PCD through a different pathway. Among the target genes regulated by the redox rhythm, both EDS1 and PAD4 (C2) are critical regulators of PCD and resistance during ETI, even though they are not required for the function of the coiled-coil (CC) class of NB-LRRs^39^ used in this study. Furthermore, C2 and C6 also include regulators of immune responses mediated by two other plant defence hormones, jasmonic acid (JA) and ethylene (ET). GSNOR1 is known to both regulate PCD^23, 40^ and promote JA-induced defence against herbivores in tobacco^41^. SOBIR1 and RECEPTOR LIKE PROTEINs (RLPs, in C2) are involved in PCD^42, 43^ and JA/ET-mediated defence against necrotrophic fungi^44^. Moreover, a major role for EDS1/PAD4 in promoting PCD is through suppression of the MYC2 transcription factor (TF) in the JA pathway^45^. Combined with the presence of the JA receptor CORONATINE INSENSITIVE 1 (COI1) in C2 and the requirement for JA production and signalling for full induction of PCD^46^, these findings suggest that the circadian redox rhythm may regulate PCD through the JA and/or the ET defence pathways.

The JA defence pathway is normally suppressed by the JASMONATE-ZIM-DOMAIN PROTEIN (JAZ) proteins which are degraded during ETI^46^. Downstream of JAZ, the JA pathway contains two mutually antagonistic branches: the JA-only branch with MYC2 as the central TF promoting defence against herbivores and the shared JA/ET branch protecting against necrotrophic pathogens with ETHYLENE-INSENSITIVE3 (EIN3) and ETHYLENE-INSENSITIVE3-LIKE 1 (EIL1) as key TFs^47^.

To determine which of the JA signalling branches is involved in the circadian control of PCD, we used the non-inducible *jaz1ϕ..jas* mutant expressing a JAZ1 protein lacking the Jas degradation domain^48^ and key TF mutants *myc2*^49^ and *ein3 eil1*^50^ to assess PCD after infiltration with *Psm* ES4326/*avrRpt2*. We found that while both the *jaz1ϕ..jas* and *ein3 eil1* mutants lost their time-of-day sensitivity to ETI-triggered PCD, *myc2* was indistinguishable from WT (Fig. 4a, b), indicating that an intact JA/ET branch is necessary for circadian regulation of ETI-associated PCD.

**Fig. 4.**
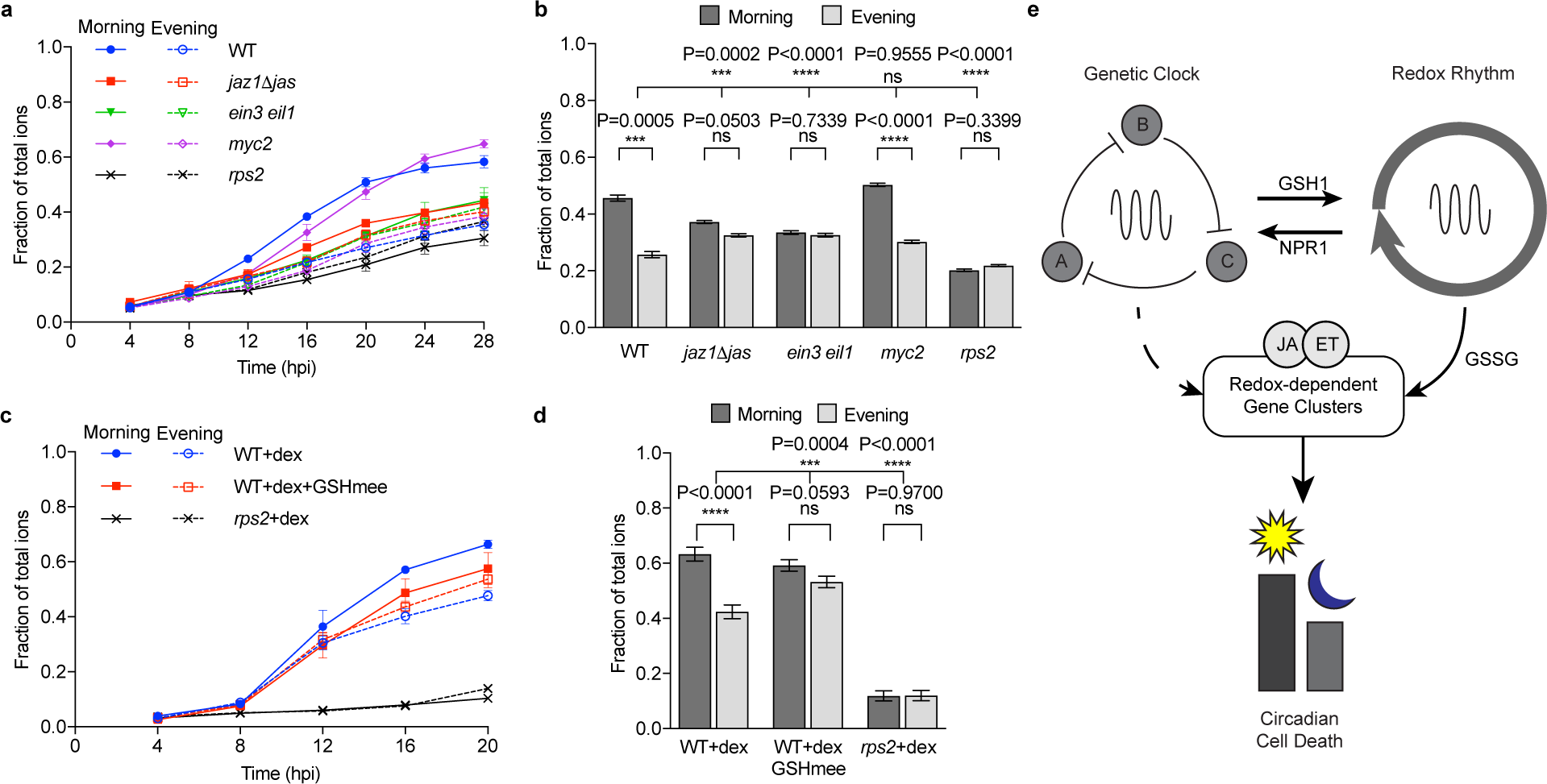
Redox rhythm regulates PCD through the JA/ET defence hormone pathway. **a**, Time-course of ion leakage after infiltration with *Psm* ES4326/*AvrRpt2* in WT, *jaz111jas*, *ein3 eil1*, *myc2*, and *rps2*. Conductivity was normalized to total ions for each sample. n =3. **b**, Analysis of normalized conductivity at 20 hpi from three separate experiments combined using a linear mixed-effect model. n = 9. Individual values were compared using Student’s t-test, while the interaction with the WT was tested using 2-way ANOVA; ns, not significant. **c**, Time-course of ion leakage of WT plants carrying *Dex:AvrRpt2* after treatment with dex or dex + GSHmee, and *rps2* plants expressing *Dex:AvrRpt2* after treatment with dex. n = 3. **d**, Analysis of normalized conductivity at 20 hpi from three separate experiments was carried out as in (**b**). All values are means ± SEM. **e**, The model of circadian regulation of immune-induced PCD in plants. Genetic clock and redox rhythm are coupled through proteins such as NPR1 and GSH1. Redox rhythm has distinct transcriptional targets and promotes PCD in the morning through the jasmonic acid/ethylene (JA/ET) defence pathway.

Finally, to further demonstrate the role of the redox rhythm in the circadian regulation of PCD, we used the cell permeable GSH derivative, glutathione monoethyl ester (GSHmee), to perturb the cellular redox rhythm. GSHmee treatment has previously been shown to disrupt the redox rhythm while reinforcing the genetic clock through the activity of NPR1^14^. To exclude a possible influence of GSHmee on the growth of the pathogen, we used the transgenic plants that express the bacterial effector avrRpt2 in response to dexamethasone (dex) treatment to induce PCD. As expected, in the absence of the bacterial carrier, *in planta* expression of the effector through dex induction alone is sufficient to induce PCD. More importantly, a stronger PCD was observed in the subjective morning than in the subjective evening, while no difference was observed after dex and GSHmee combined treatment (Fig. 4c, d). This further links cellular glutathione levels to time-of-day sensitivity to ETI induction of PCD and demonstrates that this process is independent of the bacterial pathogen.

Together, our study clearly establishes that ETI-triggered PCD is a biological process specifically regulated by the circadian redox rhythm through the JA/ET defence signalling pathway (Fig. 4e). This circadian control of PCD is distinct from the genetic-clock-regulated expression of antimicrobial genes mediated by the defence hormone SA and NPR1 in promoting cell survival and resistance in response to pathogen challenge^14, 38^. Despite of their distinguishable roles in regulating different plant immune responses, redox rhythm and genetic clock are intrinsically intertwined through the activities of proteins such as NPR1 which responds to redox perturbations by enhancing the amplitude of the genetic clock^14^. In the reverse direction, the genetic clock controls the production of redox molecules, such as glutathione, through transcription of genes, such as *GSH1* (Extended Data Fig. 1c, d, Fig. 4e). The use of the long-period mutant *prr7 prr9* allowed us to disentangle the two oscillatory systems, at least transiently, under LL. This dynamic interplay between the two oscillators in *prr7 prr9* might be the cause for our inability to obtain a consistent PCD phenotype in this mutant.

The regulation of ETI-associated PCD by the redox oscillation, rather than the genetic clock, might be due to the intensive energy requirement for this immune response which is known to involve the expenditure of ATP^6, 31^ and transcriptional and translational reprogramming^51^. Gating PCD towards morning would allow the response to coincide with the production peaks of energy (ATP)^52^ and reducing power (NADPH^14^and GSH)(Extended Data Fig.1a). Moreover, the circadian oscillations of basal JA, whose synthesis involves beta oxidation^47^, also peak in mid morning^53^, suggesting a possible link between the redox rhythm and JA production. Similar to the defence response, flowering is not only impacted by GSH levels^54^, but also requires sufficient energy reserves (i.e., high carbohydrate content), evident from the requirement of a fully functional trehalose-6-phosphate pathway for *FT* expression and oscillation^55^. Based on the results of this study, we hypothesize that with redox rhythm serving as a sensitive signalling hub, organisms gain greater flexibility in minimizing metabolic conflict caused by pathogen challenge or transition from growth to reproduction.

The scope of this study was defined by the limited pool of the probes used in the RASL-seq of *prr7 prr9* and the measurement of a single redox pair of GSH/GSSG due to our research focus and technical limitations. The incidental detection of the flowering time gene *FT* which oscillates with a shorter period in the long period *prr7 prr9* mutant even at 30 °C, indicates the presence of other target genes and physiological processes that are controlled by the redox rhythm. A more comprehensive investigation involving time-course measurements of the whole transcriptome and metabolome using mutants in which both genetic and redox oscillators are present, but with different period lengths, would allow the expansion of the study beyond ETI-mediated PCD and reveal a fuller picture of the role that the redox rhythm plays in regulating plant physiology. The conservation of this ancient oscillator across all lineages of life^3^ suggests that regulation of major incidental energy-consuming events, such as immune-induced PCD and flowering, might be the universal reason for retaining the circadian redox oscillator during evolution even after the subsequent emergence of the genetic clock.

## Acknowledgements

We thank Dr. George H. Greene for designing the probe set for RASL-seq, Dr. Tianyuan Chen for assisting in preparation of the RASL-seq library for sequencing, Ethan Gurwitch for assistance with plant maintenance and Dr. Yucong Xie for help with statistical analysis. We also thank Dr. Zhonglin Mou for gifting *pgl3* and *pgl3 npr1* seeds, Dr. Hongwei Guo for *ein3 eil1* and *myc2* seeds, Dr. Xing Zhang for helpful discussion on the project and Dr. Steve B. Hasse for critical reading of the manuscript.

## Author contributions

S.K. and X.D. designed the study. S.K. performed the experiments, data processing and statistical analysis with M.M. helping with tissue collection, preparation of the RASL-seq library for sequencing and initial data processing for RASL-seq. S.K. and X.D. prepared the manuscript. All the authors discussed and revised the manuscript.

## Data and materials availability

Source data will be available upon acceptance. Requests for materials, reagents, and scripts will be addressed to the corresponding author Xinnian Dong.

---

**Extended data Fig. 1.**
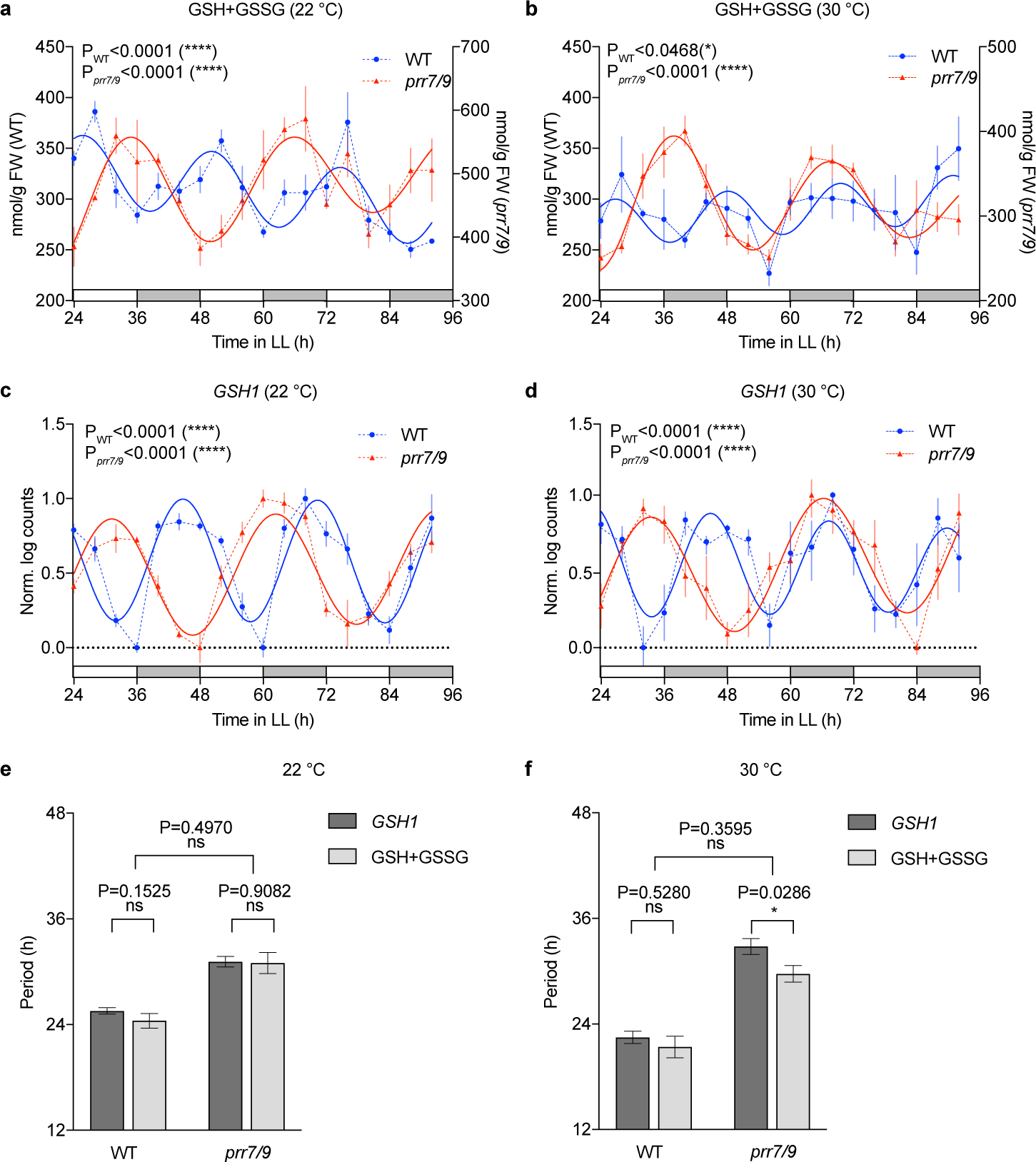
Total glutathione oscillation is driven by the genetic clock through regulation of *GSH1*. **a**, **b**, Total glutathione levels in WT and *prr7 prr9* at 22 °C (**a**) and 30 °C (**b**). Solid lines represent the harmonic regression, indicating statistically significant oscillations. n = 3 for (**a**); n ≥ 4 for (**b**). **c**, **d**, Normalized *GSH1* transcript levels in WT and *prr7 prr9* at 22 °C (**c**) and 30 °C (**d**). n ≥ 5 for (**c**); n = 3 for (**d**). Solid lines represent the harmonic regression, indicating statistically significant oscillations. n ≥ 5 for (**c**); n = 3 for (**d**). **e** and **f**, Estimated periods from the harmonic regression for (**a**-**d**). Individual values were compared using Student’s t-test, while the interaction was tested using 2-way ANOVA; ns, not significant. All values are means ± SEM.

**Extended data Fig. 2.**
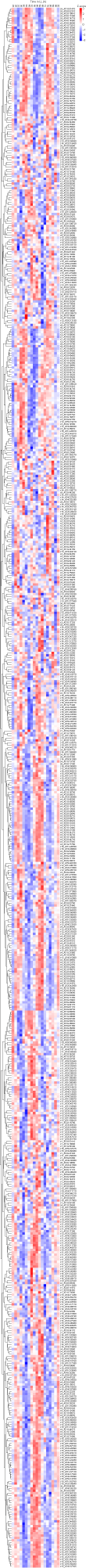
Deterministic clustering supports the results from k-means clustering. The heatmap shows the hierarchical clustering of the RASL-seq dataset from the *prr7 prr9* at 22 °C samples based on Euclidean distance using complete linkage. Zoom to see individual transcripts. The columns represent the timepoints while the rows are labelled with the gene model identifier preceded with the cluster number assigned by the k-means clustering analysis.

**Extended data Fig. 3.**
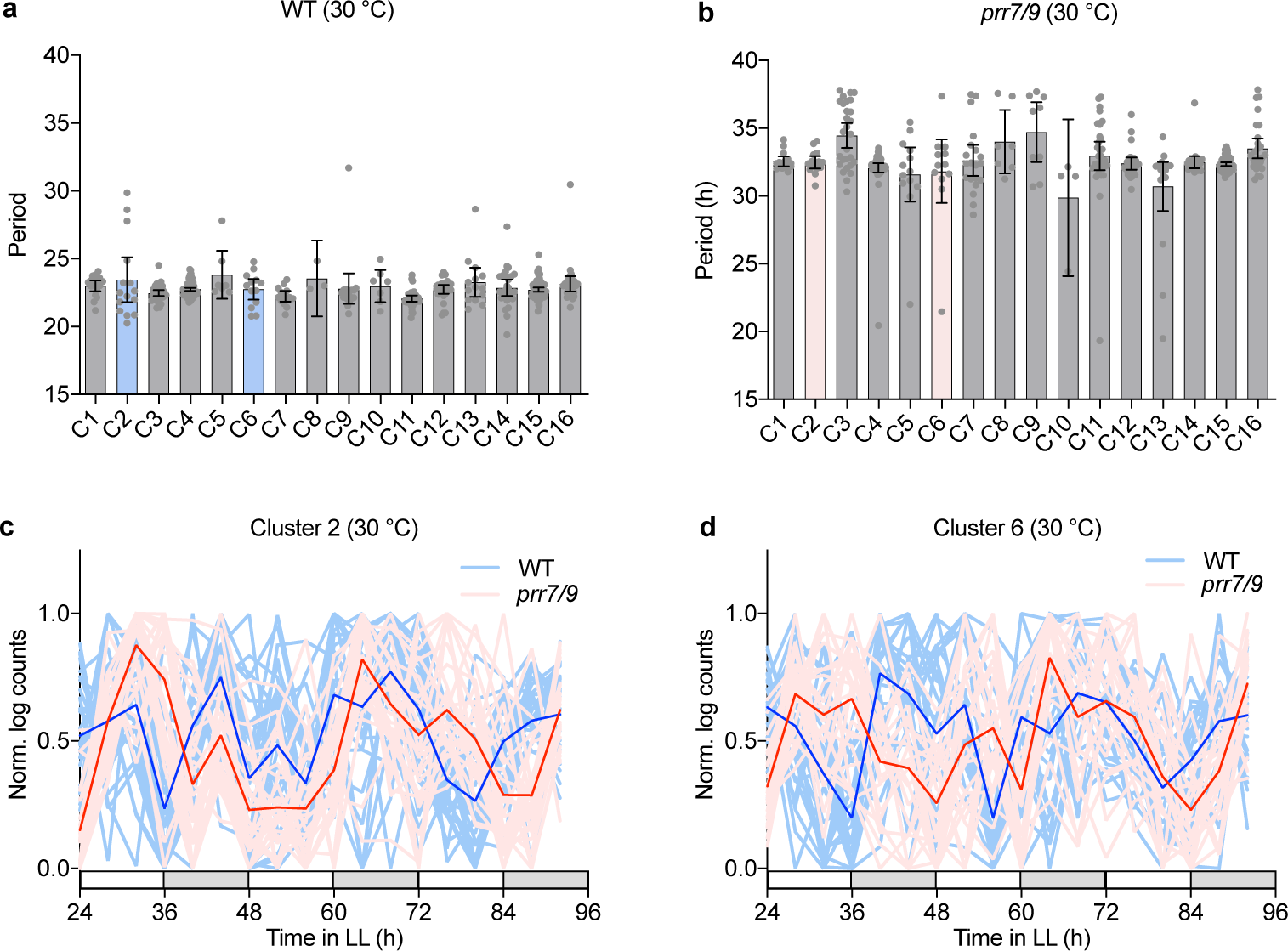
Genetic clock is the predominant regulator of candidate gene clusters at higher temperatures. **a**, **b**, Periods for oscillatory transcripts in WT (**a**) and in *prr7 prr9* (**b**) at 30 °C. Clusters were generated based on k-means analysis of expression signatures of all transcripts in the RASL-seq dataset from the *prr7 prr9* at 22 °C samples. Periods were estimated using FFT-NLLS, with each dot representing the period of a single transcript. Clusters of interest, C2 and C6, are coloured light blue (**a**) and light red (**b**), respectively. **c**, **d**, Normalized expression levels of all genes in C2 (**c**) and C6 (**d**) in WT and *prr7 prr9* at 30 °C. Dark lines represent the means of each cluster, while lighter lines represent the expression pattern of the means of individual genes in the clusters.

**Extended data Fig. 4.**
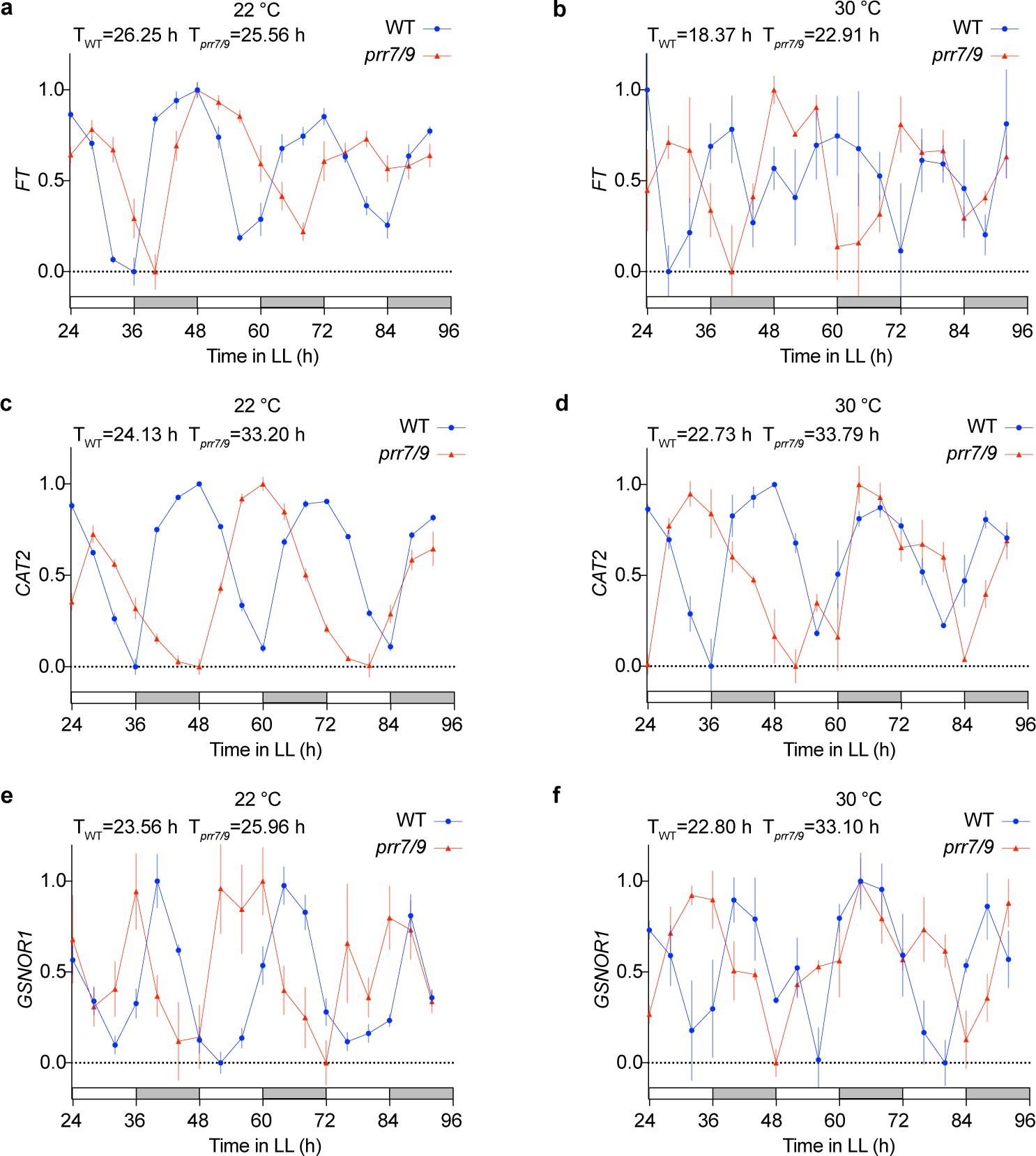
Higher temperature delinks *GSNOR1,* but not *FT*, from the redox rhythm. **a**-**f**, Normalized expressions of *FT* (**a**, **b**), *CAT2* (**c**, **d**), and *GSNOR1* (**e**, **f**) from RASL-seq at 22 °C (left) and 30 °C (right). Values are means ± SEM. n ≥ 5 for 22 °C; n = 3 for 30 °C. The periods were determined using harmonic regression.

**Extended data Fig. 5.**
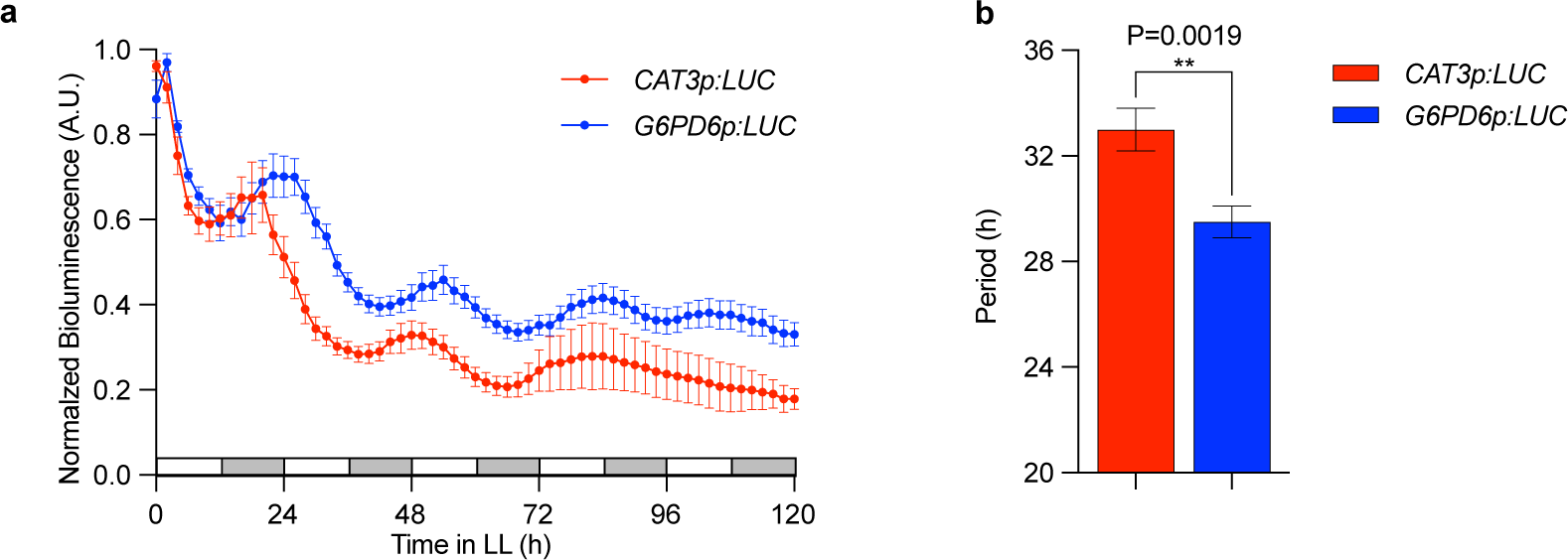
Luminescence imaging confirms that *G6PD6* from cluster 2 oscillates with a shorter period than the evening-phased clock output *CAT3*. **a,** Normalized bioluminescence traces for independent T1 transformants of *CAT3p:LUC* and *G6PD6p:LUC* with n = 10 and n = 13, respectively, in *prr7 prr9*. **b,** Period estimation (FFT-NLLS) of independent T1 transformants from two separate experiments, combined using a linear mixed-effect model, for *CAT3p:LUC* and *G6PD6p:LUC* in *prr7 prr9* with n = 10 and n = 19, respectively. Only oscillations with RAE ≤ 0.7 were considered for the analysis. Periods were compared using Student’s t-test; All values are means ± SEM.

## References

1 Dodd, A. N. et al. Plant Circadian Clocks Increase Photosynthesis, Growth, Survival, and Competitive Advantage. Science 309, 630–633 (2005).

2 O’Neill, J. S. & Reddy, A. B. Circadian clocks in human red blood cells. Nature 469, 498–503 (2011). https://doi.org:http://www.nature.com/nature/journal/v469/n7331/abs/10.1038-nature09702-unlocked.html#supplementary-information

3 Edgar, R. S. et al. Peroxiredoxins are conserved markers of circadian rhythms. Nature 485, 459–464 (2012). https://doi.org:http://www.nature.com/nature/journal/v485/n7399/abs/nature11088.html#supplementary-information

4 Abruzzi, K. C., Gobet, C., Naef, F. & Rosbash, M. Comment on “Circadian rhythms in the absence of the clock gene Bmal1”. Science 372, eabf0922 (2021). https://doi.org:10.1126/science.abf0922

5 Salomé, P. A., Weigel, D. & McClung, C. R. The role of the Arabidopsis morning loop components CCA1, LHY, PRR7, and PRR9 in temperature compensation. Plant Cell 22, 3650-3661 (2010). https://doi.org:10.1105/tpc.110.079087

6 Chen, T. et al. Global translational induction during NLR-mediated immunity in plants is dynamically regulated by CDC123, an ATP-sensitive protein. Cell Host & Microbe (2023). https://doi.org/10.1016/j.chom.2023.01.014

7 >McClung, C. R. Plant circadian rhythms. Plant Cell 18, 792–803 (2006). https://doi.org:10.1105/tpc.106.040980

8 Bargiello, T. A. & Young, M. W. Molecular genetics of a biological clock in Drosophila. Proc Natl Acad Sci U S A 81, 2142–2146 (1984). https://doi.org:10.1073/pnas.81.7.2142

9 Zehring, W. A. et al. P-element transformation with period locus DNA restores rhythmicity to mutant, arrhythmic Drosophila melanogaster. Cell 39, 369–376 (1984). https://doi.org:10.1016/0092-8674(84)90015-1

10 Vitaterna, M. H. et al. Mutagenesis and mapping of a mouse gene, Clock, essential for circadian behavior. Science 264, 719–725 (1994). https://doi.org:10.1126/science.8171325

11 Greenham, K. & McClung, C. R. Integrating circadian dynamics with physiological processes in plants. Nature Reviews Genetics 16, 598 (2015). https://doi.org:10.1038/nrg3976

12 Rey, G. et al. Metabolic oscillations on the circadian time scale in Drosophila cells lacking clock genes. Molecular Systems Biology 14, e8376 (2018). https://doi.org/10.15252/msb.20188376

13 Ray, S. et al. Circadian rhythms in the absence of the clock gene Bmal1. Science 367, 800–806 (2020). https://doi.org:10.1126/science.aaw7365

14 Zhou, M. et al. Redox rhythm reinforces the circadian clock to gate immune response. Nature 523, 472–476 (2015). https://doi.org:10.1038/nature14449

15 Rey, G. et al. The Pentose Phosphate Pathway Regulates the Circadian Clock. Cell Metabolism 24, 462–473 (2016). https://doi.org:10.1016/j.cmet.2016.07.024

16 Putker, M. et al. Mammalian Circadian Period, But Not Phase and Amplitude, Is Robust Against Redox and Metabolic Perturbations. Antioxid Redox Signal (2017). https://doi.org:10.1089/ars.2016.6911

17 Mwimba, M. et al. Daily humidity oscillation regulates the circadian clock to influence plant physiology. Nature Communications 9, 4290 (2018). https://doi.org:10.1038/s41467-018-06692-2

18 Foyer, C. H. & Noctor, G. Ascorbate and glutathione: the heart of the redox hub. Plant Physiol 155, 2–18 (2011). https://doi.org:10.1104/pp.110.167569

19 Yeakley, J. M. et al. Profiling alternative splicing on fiber-optic arrays. Nature Biotechnology 20, 353–358 (2002). https://doi.org:10.1038/nbt0402-353

20 Pittendrigh, C. S. ON TEMPERATURE INDEPENDENCE IN THE CLOCK SYSTEM CONTROLLING EMERGENCE TIME IN DROSOPHILA*†. Proceedings of the National Academy of Sciences 40, 1018–1029 (1954). https://doi.org:10.1073/pnas.40.10.1018

21 Xiang, C., Werner, B. L., Christensen, E. M. & Oliver, D. J. The biological functions of glutathione revisited in arabidopsis transgenic plants with altered glutathione levels. Plant Physiol 126, 564–574 (2001). https://doi.org:10.1104/pp.126.2.564

22 Zielinski, T., Moore, A. M., Troup, E., Halliday, K. J. & Millar, A. J. Strengths and Limitations of Period Estimation Methods for Circadian Data. PLOS ONE 9, e96462 (2014). https://doi.org:10.1371/journal.pone.0096462

23 Feechan, A. et al. A central role for S-nitrosothiols in plant disease resistance. Proceedings of the National Academy of Sciences 102, 8054–8059 (2005). https://doi.org:10.1073/pnas.0501456102

24 Liu, L. et al. A metabolic enzyme for S-nitrosothiol conserved from bacteria to humans. Nature 410, 490–494 (2001). https://doi.org:10.1038/35068596

25 Kim, H., Kim, Y., Yeom, M., Lim, J. & Nam, H. G. Age-associated circadian period changes in Arabidopsis leaves. J Exp Bot 67, 2665–2673 (2016). https://doi.org:10.1093/jxb/erw097

26 Takahashi, N., Hirata, Y., Aihara, K. & Mas, P. A Hierarchical Multi-oscillator Network Orchestrates the Arabidopsis Circadian System. Cell 163, 148–159 (2015). https://doi.org/10.1016/j.cell.2015.08.062

27 Wakao, S. & Benning, C. Genome-wide analysis of glucose-6-phosphate dehydrogenases in Arabidopsis. The Plant Journal 41, 243–256 (2005). https://doi.org/10.1111/j.1365-313X.2004.02293.x

28 Gao, M. et al. Regulation of Cell Death and Innate Immunity by Two Receptor-like Kinases in Arabidopsis. Cell Host & Microbe 6, 34–44 (2009). https://doi.org:10.1016/j.chom.2009.05.019

29 Rustérucci, C., Aviv, D. H., Holt, B. F., Dangl, J. L. & Parker, J. E. The Disease Resistance Signaling Components EDS1 and PAD4 Are Essential Regulators of the Cell Death Pathway Controlled by LSD1 in Arabidopsis. The Plant Cell 13, 2211–2224 (2001). https://doi.org:10.1105/tpc.010085

30 Torres, M. A., Dangl, J. L. & Jones, J. D. G. Arabidopsis gp91phox homologues AtrbohD and AtrbohF are required for accumulation of reactive oxygen intermediates in the plant defense response. Proceedings of the National Academy of Sciences 99, 517–522 (2002). https://doi.org:10.1073/pnas.012452499

31 Hatsugai, N., Perez Koldenkova, V., Imamura, H., Noji, H. & Nagai, T. Changes in cytosolic ATP levels and intracellular morphology during bacteria-induced hypersensitive cell death as revealed by real-time fluorescence microscopy imaging. Plant Cell Physiol 53, 1768–1775 (2012). https://doi.org:10.1093/pcp/pcs119

32 Korneli, C., Danisman, S. & Staiger, D. Differential Control of Pre-Invasive and Post-Invasive Antibacterial Defense by the Arabidopsis Circadian Clock. Plant and Cell Physiology 55, 1613–1622 (2014). https://doi.org:10.1093/pcp/pcu092

33 Menna, A., Nguyen, D., Guttman, D. S. & Desveaux, D. Elevated Temperature Differentially Influences Effector-Triggered Immunity Outputs in Arabidopsis. Front Plant Sci 6, 995 (2015). https://doi.org:10.3389/fpls.2015.00995

34 Wang, Z.-Y. & Tobin, E. M. Constitutive Expression of the *CIRCADIAN CLOCK ASSOCIATED 1* (*CCA1*) Gene Disrupts Circadian Rhythms and Suppresses Its Own Expression. Cell 93, 1207–1217 (1998). https://doi.org:10.1016/S0092-8674(00)81464-6

35 Zhang, C. et al. Crosstalk between the Circadian Clock and Innate Immunity in Arabidopsis. Plos Pathogens 9 (2013).

36 Xiong, Y., DeFraia, C., Williams, D., Zhang, X. & Mou, Z. Characterization of Arabidopsis 6-Phosphogluconolactonase T-DNA Insertion Mutants Reveals an Essential Role for the Oxidative Section of the Plastidic Pentose Phosphate Pathway in Plant Growth and Development. Plant and Cell Physiology 50, 1277–1291 (2009). https://doi.org:10.1093/pcp/pcp070

37 Ch, R. et al. Rhythmic glucose metabolism regulates the redox circadian clockwork in human red blood cells. Nature Communications 12, 377 (2021). https://doi.org:10.1038/s41467-020-20479-4

38 Zavaliev, R., Mohan, R., Chen, T. & Dong, X. Formation of NPR1 Condensates Promotes Cell Survival during the Plant Immune Response. Cell 182, 1093–1108.e1018 (2020). https://doi.org:10.1016/j.cell.2020.07.016

39 Aarts, N. et al. Different requirements for EDS1 and NDR1 by disease resistance genes define at least two R gene-mediated signaling pathways in Arabidopsis. Proceedings of the National Academy of Sciences 95, 10306–10311 (1998). https://doi.org:10.1073/pnas.95.17.10306

40 Yun, B.-W. et al. S-nitrosylation of NADPH oxidase regulates cell death in plant immunity. Nature, 264–268 (2011). https://doi.org:http://www.nature.com/nature/journal/v478/n7368/abs/nature10427.html#supplementary-information

41 Wunsche, H., Baldwin, I. T. & Wu, J. S-Nitrosoglutathione reductase (GSNOR) mediates the biosynthesis of jasmonic acid and ethylene induced by feeding of the insect herbivore Manduca sexta and is important for jasmonate-elicited responses in Nicotiana attenuata. J Exp Bot 62, 4605–4616 (2011). https://doi.org:10.1093/jxb/err171

42 Liebrand, T. W. H. et al. Receptor-like kinase SOBIR1/EVR interacts with receptor-like proteins in plant immunity against fungal infection. Proceedings of the National Academy of Sciences 110, 10010–10015 (2013). https://doi.org:10.1073/pnas.1220015110

43 Ma, L. & Borhan, M. H. The receptor-like kinase SOBIR1 interacts with Brassica napus LepR3 and is required for Leptosphaeria maculans AvrLm1-triggered immunity. Front Plant Sci 6 (2015). https://doi.org:10.3389/fpls.2015.00933

44 Zhang, W., et al. *Arabidopsis* RECEPTOR-LIKE PROTEIN30 and Receptor-Like Kinase SUPPRESSOR OF BIR1-1/EVERSHED Mediate Innate Immunity to Necrotrophic Fungi. The Plant Cell 25, 4227–4241 (2013). https://doi.org:10.1105/tpc.113.117010

45 Cui, H. et al. Antagonism of Transcription Factor MYC2 by EDS1/PAD4 Complexes Bolsters Salicylic Acid Defense in Arabidopsis Effector-Triggered Immunity. Mol Plant 11, 1053–1066 (2018). https://doi.org/10.1016/j.molp.2018.05.007

46 Liu, L. et al. Salicylic acid receptors activate jasmonic acid signalling through a non-canonical pathway to promote effector-triggered immunity. Nat Commun 7, 13099 (2016). https://doi.org:10.1038/ncomms13099 https://www.nature.com/articles/ncomms13099#supplementary-information

47 Wang, Y., Mostafa, S., Zeng, W. & Jin, B. Function and Mechanism of Jasmonic Acid in Plant Responses to Abiotic and Biotic Stresses. Int J Mol Sci 22 (2021). https://doi.org:10.3390/ijms22168568

48 Thines, B. et al. JAZ repressor proteins are targets of the SCF(COI1) complex during jasmonate signalling. Nature 448, 661–665 (2007). https://doi.org:10.1038/nature05960

49 Lorenzo, O., Chico, J. M., Sánchez-Serrano, J. J. & Solano, R. JASMONATE-INSENSITIVE1 encodes a MYC transcription factor essential to discriminate between different jasmonate-regulated defense responses in Arabidopsis. Plant Cell 16, 1938–1950 (2004). https://doi.org:10.1105/tpc.022319

50 Alonso, J. M. et al. Five components of the ethylene-response pathway identified in a screen for weak ethylene-insensitive mutants in Arabidopsis. Proc Natl Acad Sci U S A 100, 2992–2997 (2003). https://doi.org:10.1073/pnas.0438070100

51 Yoo, H. et al. Translational Regulation of Metabolic Dynamics during Effector-Triggered Immunity. Mol Plant 13, 88–98 (2020). https://doi.org:10.1016/j.molp.2019.09.009

52 Cervela-Cardona, L. et al. Circadian Control of Metabolism by the Clock Component TOC1. Front Plant Sci 12, 683516 (2021). https://doi.org:10.3389/fpls.2021.683516

53 Goodspeed, D., Chehab, E. W., Min-Venditti, A., Braam, J. & Covington, M. F. Arabidopsis synchronizes jasmonate-mediated defense with insect circadian behavior. Proceedings of the National Academy of Sciences 109, 4674–4677 (2012). https://doi.org:10.1073/pnas.1116368109

54 Ogawa, K. i., Tasaka, Y., Mino, M., Tanaka, Y. & Iwabuchi, M. Association of Glutathione with Flowering in Arabidopsisthaliana. Plant and Cell Physiology 42, 524–530 (2001). https://doi.org:10.1093/pcp/pce065

55 Wahl, V. et al. Regulation of Flowering by Trehalose-6-Phosphate Signaling in Arabidopsis thaliana. Science 339, 704–707 (2013).

